# Beacon Reconstruction Attack: Reconstruction of genomes in genomic data-sharing beacons using summary statistics

**DOI:** 10.1101/2024.12.10.627379

**Authors:** Kousar Saleem, A. Ercument Cicek, Sinem Sav

## Abstract

Genomic data sharing beacon protocol, developed by the Global Alliance for Genomics and Health (GA4GH), offers a privacy-preserving mechanism for querying genomic datasets while restricting direct data access. Despite their design, beacons remain vulnerable to privacy attacks. This study introduces a novel privacy vulnerability of the protocol: One can reconstruct large portions of the genomes of all beacon participants by only using the summary statistics reported by the protocol. We introduce a novel optimization-based algorithm that leverages beacon responses and single nucleotide polymorphism (SNP) correlations for reconstruction. By optimizing for the SNP correlations and allele frequencies, the proposed approach achieves genome reconstruction with a substantially higher F1-score (70%) compared to baseline methods (45%) on beacons generated using individuals from the HapMap and OpenSNP datasets. Our findings reveal critical vulnerabilities in beacon protocol, underscoring the need for enhanced privacy-preserving mechanisms to protect genomic data. Our implementation is available at https://github.com/ASAP-Bilkent/Beacon-Reconstruction-Attack.

## Introduction

The rapid advancements in genomic research have led to unprecedented access to vast amounts of genomic data, enabling breakthroughs in precision medicine, disease prediction, and population genetics. However, sharing such sensitive data comes with significant privacy challenges (1; 2). Genomic data beacons (3), a widely adopted approach for secure genomic data sharing, aim to balance the need for data accessibility with the requirement for individual privacy. Introduced by the Global Alliance for Genomics and Health (GA4GH) (4), beacons provide an interface for the researchers to query datasets to ask whether a specific allele at a specific position exists in the dataset. The protocol responds with a simple “yes/no” answer. If the answer is “yes”, it can also report the frequency of the allele in the underlying datasets (5). This mechanism enables collaboration without direct access to the data itself, reducing the risks of exposing sensitive genomic information.

Genomic beacons are not immune to privacy attacks, despite being designed to enable secure querying of genomic databases. One of the most notable threats is the membership inference attacks, in which an adversary attempts to determine whether a particular individual’s genomic data is in the queried dataset. The adversary can accumulate information through subsequent queries, exploiting statistical patterns to make inferences about the composition of the dataset. This presents substantial privacy challenges, particularly because beacons are often linked to specific sensitive phenotypes such as the Autism Speaks MSSNG Dataset (6). Several works have introduced approaches based on likelihood ratio test to launch membership inference attacks against beacons (7; 8; 9). Several countermeasures have been proposed against these attacks with techniques including flipping responses (10), query budgets for users (8), including relatives of the participants in the databases (11), differential privacy (12), game theory (13; 14; 15) and reinforcement learning (15). In addition to membership inference attacks, beacons are also vulnerable to *genome reconstruction attacks* as first introduced by Ayoz et al. (16). This method leverages SNP correlations along with clustering techniques to reconstruct large portions of individual genomes using the responses of the beacon system. The attack exploits the dynamic nature of a beacon in which participants are added/removed over time, and the changes in responses must belong to the added/removed beacon participants (16).

In this study, we show that the summary statistics reported by the beacon reveal more than expected by design about the genomes within the database and we introduce a new genome reconstruction attack. For the first time, we demonstrate that an adversary can reconstruct genomes of all individuals in a beacon database by using only the beacon responses. That is, an adversary who takes the snapshot of the beacon (e.g., query for a target SNP subset of interest) can use the correlations among SNPs to learn which participants carry each SNPs. To achieve this, we introduce a two-step optimization-based algorithm that optimizes the following objectives in alternating fashion: (i) Matching the publicly available correlations among the SNPs, and (ii) Matching the frequencies reported by the beacon. Our results on beacons generated using HapMap and OpenSNP datasets demonstrate that given a randomly selected set of 1000 SNPs and a beacon with 50 individuals, an adversary can reconstruct all genomes with an F1-score of 70% while the baseline approach can only achieve 45%. We analyze the effect of increasing number of individuals and SNPs. We further show that the attack becomes stronger if the adversary has access to the genomes of a portion of the people in the beacon and the attack is feasible even if the beacon does not report allele frequencies. Our implementation is available at https://github.com/ASAP-Bilkent/Beacon-Reconstruction-Attack.git.

## Related Work

Numerous studies have shown that anonymization of genomic samples deos not effectively preserve the privacy of the individuals. Homer et al. demonstrated that individuals contributing even trace amounts of DNA to Genome-Wide Association Studies (GWAS) with complex mixtures can be reidentified (17). Wang et al. demonstrated that information leaks from GWAS can lead to the identification of individuals through linkage to public datasets (18). Gymrek et al. further exposed vulnerabilities in anonymized genomic data by demonstrating how surname inference from Y-chromosome can be combined with publicly available genealogical databases to re-identify participants in genomic studies (19). Similarly, Erlich et al. used linkage attacks by integrating anonymized genomic data with auxiliary information from genealogical sources, uncovering new privacy risks (20). Humbert et al. and Samani et al. examined the potential of using high-order SNP correlations and familial ties to infer hidden genomic information (21; 22).

In response to the limitations of traditional anonymization techniques, Differential Privacy (DP) emerged as an advanced approach to privacy preservation. DP ensures that the inclusion or exclusion of an individual’s data in a dataset minimally affects the results of any analysis, thus safeguarding personal privacy. Initially introduced by Dwork et al. through the concept of adding noise proportional to query sensitivity (23; 24). This framework has been widely applied in Genome-Wide Association Studies (GWAS) to protect sensitive genetic information (25; 26; 27). However, while DP offers robust privacy protection, it inherently involves a trade-off between privacy and data utility. To address privacy concerns while retaining data utility, genomic beacons were introduced by the Global Alliance for Genomics and Health (GA4GH) (4).

Beacons offer summary statistics to user queries about the existence of certain alleles at certain positions. These summary statistics could be a binary response or the frequency of the allele (5). Thus, the system does not reveal the entire dataset content. It provides a more controlled and secure method of data sharing as the data is shared only if the allele of interest is present in the dataset. Moreover, the summary statistics in theory do not reveal the genome or the identity of the participants. However, beacons remain vulnerable to certain attacks. Shringarpure and Bustamante demonstrated that attackers could exploit statistical techniques, such as the Likelihood Ratio Test (LRT), to infer whether an individual is part of a beacon dataset by analyzing the yes/no responses to a query set (7). This work marked a critical advancement in highlighting privacy risks, especially in scenarios where genomic data is linked to sensitive phenotypic traits. Raisaro et al. further showed that targeting SNPs with Minor Allele Frequencies (MAFs) can dramatically reduce the number of queries required for re-identification (8). Finally, von Thenen et al. used the correlations among SNPs to infer the answers of the beacon to queries that are not posed which enabled the attack to work with a smaller number of queries and also work even if the victim has not revealed certain high-risk SNPs to the beacon (28).

Beacon protocol is also vulnerable to the genome reconstruction attacks. However, these are relatively under-explored. In this type, the adversary can infer the genome sequence of the target individual(s) using the responses of the system. Ayöz et al. (16) showed, if the adversary knows that the victim has joined the study at time *t*, then they can compare the responses of the system at time *t −* 1 and *t* + 1 to conclude that the responses that have been flipped from “no” to “yes” belong to the victim. They further showed that (i) even if more than one person has joined the study it is possible to determine the victim’s alleles using clustering of the SNPs with respect to linkage disequilibrium, and also (ii) to link the public features of the victim (e.g., hair color) to the reconstructed genome using classifiers with high confidence. While this study showed that the beacons are vulnerable to genome reconstruction attacks, it relies on the strong assumption on the knowledge of the time that the victim joins the study and that the adversary is keeping snapshots of the beacon taken at multiple time points.

## Methods

In this section, we first explain the system model (Section 3.1) and then the threat model (Section 3.2). We formulate the problem in Section 3.3, and finally, we provide the details of our beacon reconstruction attack methods in Section 3.4.

### System Model

The beacon protocol processes incoming queries about the presence of specific alleles in its connected database(s), providing summary statistics for each dataset. For simplicity, we assume that the beacon is connected to a single database. The summary statistic can be a binary answer for the existence of the SNP of interest (29; 30; 31) or can be the allele frequency in the underlying database (32; 33).

However, even these simple “yes” or “no” responses can be exploited in several types of attacks, potentially leading to the exposure of sensitive information. This research aims to develop methods for reconstructing genomic data-sharing beacons based on query responses, with the potential to reveal critical information about individuals within the dataset.

This protocol functions within an authenticated online setting, meaning that the querier must log in before submitting one or more queries. Additionally, the system retains a record of previous queries submitted by the same querier.

### Threat Model

In our threat model, we consider a scenario where the querier functions as an attacker, aiming to reconstruct the dataset interfaced through the beacon protocol. We assume the attacker is a registered beacon user with access to publicly available information, such as the number of individuals in the dataset, allele frequencies (AFs) in the population, and the linkage disequilibrium of the single nucleotide polymorphisms (SNPs). As a registered user, an attacker can submit an unlimited number of queries, each disclosing whether specific alleles are present at particular genomic positions and, optionally, their respective frequencies. The attacker queries the beacon including *N* individuals, for a potentially large set of SNPs of interest (*M* ^′^) and captures a comprehensive snapshot of the dataset. Using the optimization-based approach we explain below, the attacker assigns the SNPs with the answer “Yes” or with allele frequency > 0, to *N* individuals in a one-to-many fashion such that the reported summary statistics and the correlations between the SNPs in the reconstructed genomes are matched.

### Problem Formulation

Let **B** denote the dataset which is accessed through a beacon. **B** can be represented as a binary matrix of size *N × M* where *N* denotes the number of individuals in the beacon and *M* denotes the number of cataloged SNPs in the human genome. Assume the attacker is interested in reconstructing a subset of these SNPs: *M* ^′^. The attacker queries the beacon for *M* ^′^ and obtains a response vector **b** of size |*M* ^′^|. **b** ∈ [0, 1] if the beacon responds with the AFs, or **b** ∈ {0, 1} if the beacon responds with presence/absence information for the SNPs.

The attacker has access to the following publicly available information: (i) AFs in the population which is represented as a vector **m** ∈ [0, 1] of size |*M* ^′^|; and (ii) the linkage disequilibrium of the SNPs which the attacker uses to construct a correlation matrix of size |*M* ^′^| *×* |*M* ^′^|: **C** ∈ [0, 1].

Following (16), we calculate **C** based on the Sokal-Michener similarity which is the percentage of matches between two binary vectors of equal size. In this context, we have a binary vector **s**_**j**_, for each SNP *j* in *M* ^′^ of size *N*_*p*_. *N*_*p*_ is the number of individuals in the population which we use to construct **C**. A value of one indicates that the individual is carrying the SNP, and zero otherwise. Then, the correlation between SNP *j* and SNP *k* is equal to 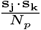

We define the problem of beacon reconstruction as an optimization problem. The goal of the attacker is to find a function *f* (**b**, *N*, **m**, *M* ^′^, **C**) to obtain a reconstruction **B**^′^ of the beacon **B** such that the Frobenius norm of their difference is minimized: 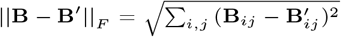.

### Beacon Reconstruction Attack

In this section, we present the first-of-its-kind beacon reconstruction attack in detail. We begin by describing our greedy (baseline) approach for beacon reconstruction, which serves as a baseline to compare our results in Section 3.4.1. Next, we introduce our novel optimization-based approach for beacon reconstruction in Section 3.4.2. Finally, we extend this optimization approach to scenarios involving known individual subsets.

### Baseline Approach for Beacon Reconstruction

In this section, we introduce a greedy algorithm in which we call the baseline approach. It is detailed in Algorithm 1.

First, the attacker initializes the matrix **B**^′^ as a zero matrix (Line 1). For each SNP *j ∈ M* ^′^ (Line 2), if the beacon responds with the frequency of the allele and the returned value is greater than zero (**b**[*j*] *>* 0) (Line 3), the attacker finds the number of individuals with this allele in the beacon simply by calculating *N* ^′^ = **b**_*j*_ *× N* (Line 4). If the beacon responses are binary, then, the attacker estimates this number using the MAF in the population which is publicly available (*N* ^′^ = **m**_*j*_ *× N*) (Line 6). The attacker then randomly selects a subset *S* of beacon participants of size *N* ^′^ that are assigned to carry the SNP *j* (Line 8).

The baseline algorithm operates under the assumption that each SNP is independent and neglects the correlation between SNPs. However, SNPs are often correlated as alleles at different loci are inherited together more than would be expected by chance because of linkage disequilibrium. To enable better reconstruction, we introduce a novel approach that uses pairwise SNP correlations in the next section.

#### Algorithm 1

Baseline Algorithm for Beacon Reconstruction Attack

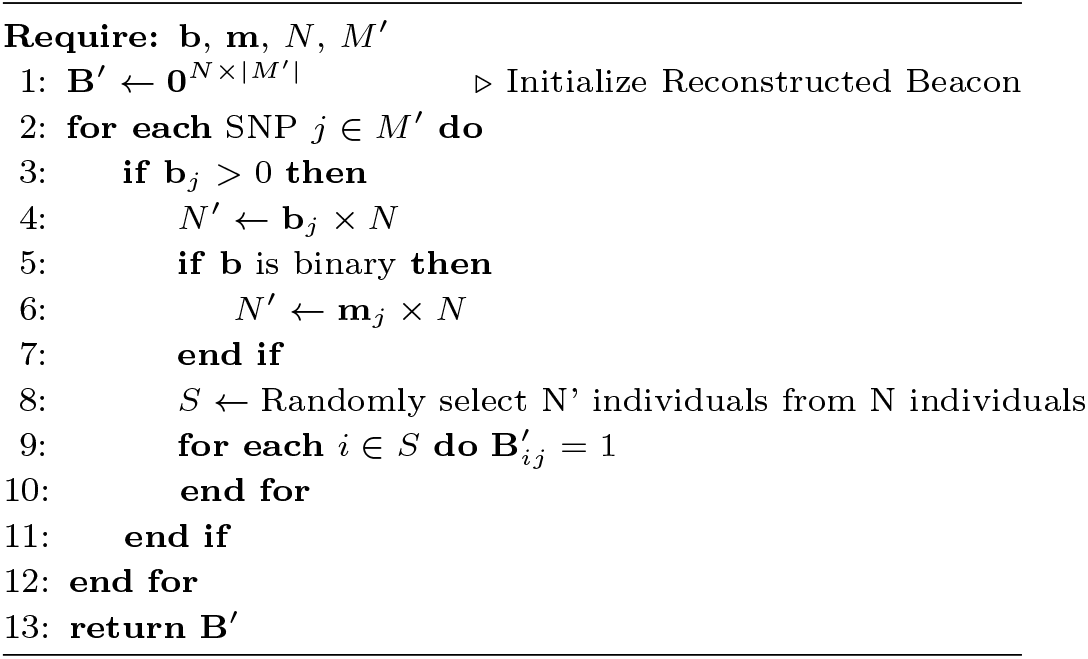

#### Optimization-based Approach for Beacon Reconstruction

The optimization-based method performs a two-objective optimization uses the pairwise SNP correlation matrix (**C**) and the allele frequencies (**b**). It iteratively refines **B**^′^ until convergence such that the reconstruction has (i) matching allele frequencies to the original beacon as reported in **b** or estimated using **m**, and (ii) matching SNP correlations to the observed SNP correlations in the population as summarized in **C**. The steps of optimization-based method are introduced in Algorithm 2 and we detail this algorithm hereafter.

##### Step 0 - Initialization

We use Algorithm 1 to initialize the reconstruction **B**^′^ based on the frequencies reported by the beacon (Line 1).

##### Step 1 - Correlation Loss Minimization

In the first step, the goal is to minimize the difference between the computed correlation matrix from **B**^′^ and the actual correlation matrix **C**, using the Frobenius norm: *L*_corr_ = 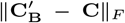 where *L*_**corr**_ is the correlation loss and 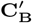 represents the correlation matrix calculated from **B**^′^. We optimize SNP assignments using the Adam optimizer (34), which updates **B**^′^ to minimize *L*_**corr**_ for *e*_1_ epochs with leaning rate *η* (Line 3 to 6). This step improves the accuracy of the reconstructed beacon by ensuring that the correlations among the SNPs are as close to the expectation. However, this might lead to mismatching allele frequencies between the actual and reconstructed beacon.

##### Step 2 - Frequency Adjustment

The second step adjusts the allele frequencies in **B**^′^ to match the original allele frequencies in **B**. That is, we want to minimize the difference between the allele frequencies obtained from the reconstruction (namely **f** ^′^), and from the actual beacon, namely **f** which is equal to **b** or if the beacon responds with binary responses, to **m**. We use the mean squared error loss: 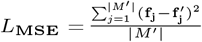. We again use the Adam optimizer, which updates **B**^′^ to minimize *L*_**MSE**_ for *e*_2_ epochs with leaning rate *η* (Line 7 to 18). We note that the is an iterative algorithm and steps 1 and 2 are repeated until convergence. Here convergence means number of flips in **B**^′^ after the update is less than a small number which we set as 10 (Line 2-19).

###### Algorithm 2

Gradient-Based Optimization Algorithm for Beacon Reconstruction Attack

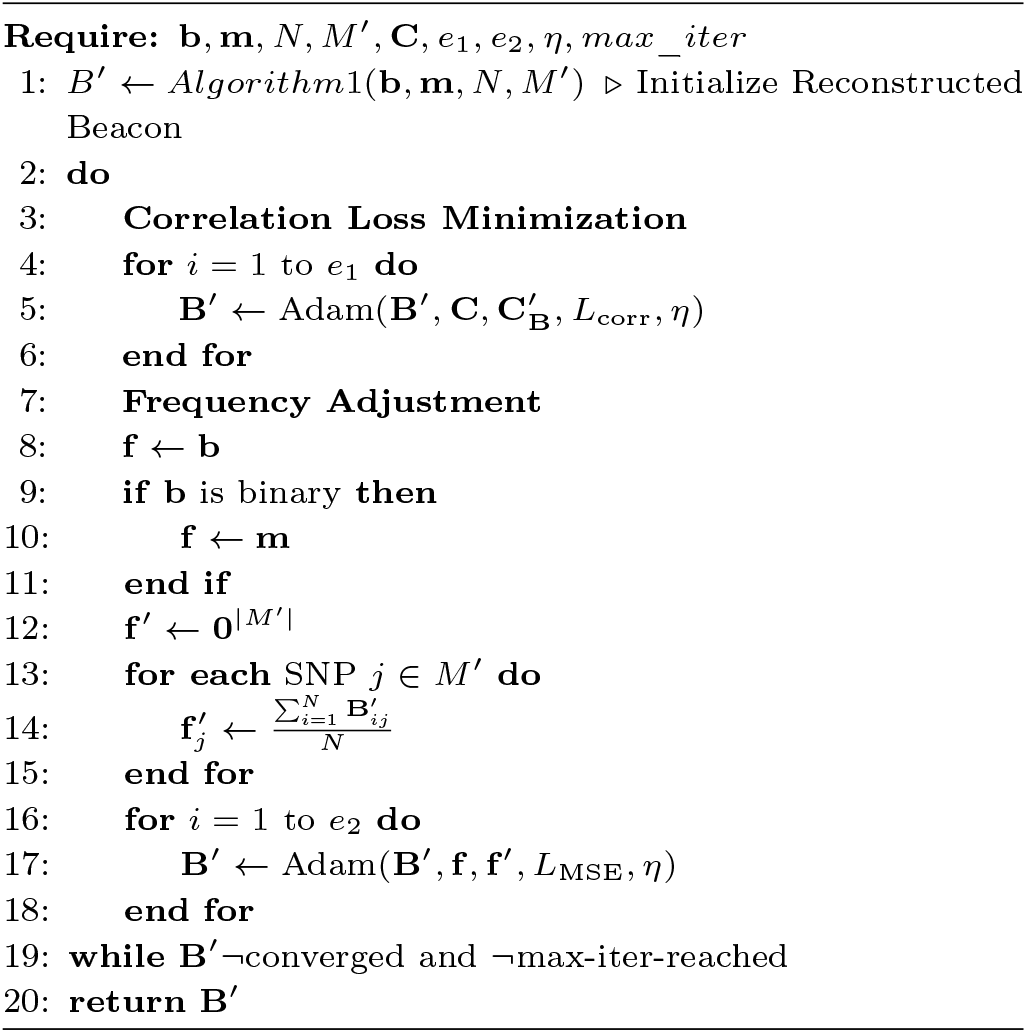

#### Reconstruction when a subset of the genomes are known by the attacker

We also consider a variant of the gradient-based optimization approach with the assumption that the attacker has access to the genomes of a percentage *p* of the beacon participants. The rationale behind this assumption is that genomic data may have already been compromised through genome reconstruction attacks targeting the beacon (16). Additionally, such data might have been accessed, either legally or illegally, from other DNA repositories, such as genealogy sites like 23andMe or ancestry.com. The attacker could then verify the membership of these individuals in the beacon using membership inference attacks (7; 8; 28).

In this scenario, the attacker aims to reconstruct the genomes of unknown individuals in the beacon, given the known individuals; first, they fix the SNP assignments of the known individuals in **B**^′^ and resonstruct the remaining individuals using Algorithm 2.

## Results

### Datasets

We use two publicly available genomic datasets to evaluate the algorithms introduced in Section 3.4: the HapMap dataset (35) and the OpenSNP dataset (36). The CEU population in the HapMap dataset contains the genotype of 164 individuals and covers over 4 million SNPs. The HapMap project is a well-established resource for understanding genetic variation, and has been previously used to evaluate attack scenarios against beacons (7; 8; 28; 16). OpenSNP contains genotypes of 2,980 individuals who donate their genomes to this community-driven project. Each individual has approximately 2 million SNPs. While the HapMap CEU population represents a beacon with a uniform genetic background, the OpenSNP dataset is diverse and due to the lack of a standard protocol for genotyping, it represents a relatively noisy and more challenging setting.

### Experimental Setup

In our experiments, we use subsets of both the HapMap and the OpenSNP datasets. We simulate beacons with *N* = 3, 10, 25, 50 and 100, where *N* represents the number of individuals in the beacon. We focus on a randomly selected set of 2000 SNPs. We experiment with various subsets *M* ^′^ of this set where |*M* ^′^| = 30, 50, 100, 500, 1000. Note that *M* ^′^ denotes the SNP subset of interest. Note that every SNP subset we construct of size |*M* ^′^|, is a superset of the smaller SNP subsets. For example, the SNP subset of size |*M* ^′^| = 100 includes the SNPs in the SNP subset of size |*M* ^′^| = 50 and smaller. We set the learning rate for the Adam optimizer as *η* = 0.001 which is applied for 1000 epochs (*e*_1_) to optimize the correlation matrix and 500 epochs (*e*_2_) to optimize the frequencies. The maximum number of iterations for the optimization algorithm to converge is set to 3000. To simulate the case where a percentage of the individuals known by the attacker, we experiment with *p* = 20% and 40%.

We construct the correlation matrix *C* using 100 left-out samples from the OpenSNP dataset to simulate the case where the publicly available correlations are relatively more noisy to test our approach in a relatively more setting. To obtain the AFs in the population, we use the 64 left-out samples from the HapMap dataset. We use the population AFs as a proxy when the beacon responds in binary instead of the exact AFs in the dataset. We also simulate the scenario where the AFs for the target population are also not available. Then, we consider using the AFs from another population. In this case, we use frequencies of the Mexican population, which we obtained from the Merged phase I+II and III HapMap Dataset^1^. All experiments are performed on a SuperMicro SuperServer with Intel Xeon CPU E5-2650 v3 2.30 GHz with 40 cores.

### Reconstruction Performance

In this subsection, we investigate the performance of the reconstuction method under varying conditions. We first focus on the effect of increasing the number of individuals in the beacon and then increasing number of targeted SNPs. Next, we investigate the performance when the attacker has already re-identified a percentage of the individuals in the beacon. Finally, we show the effect of the beacon reporting allele frequencies instead of a binary “Yes/No” answers.

#### Effect of Beacon Size

We compare the reconstruction performances of the baseline approach and the optimization-based approach with respect to the F1-score. Figure 2 shows the results of both algorithms with |*M* ^′^| = 1000, across various beacon sizes for both OpenSNP and HapMap datasets. We observe that with very few samples (*N* = 3), which is not very realistic, the baseline approach yields 64.1% and 61% F1-scores for OpenSNP and HapMap datasets, respectively. The optimization-based approach, on the other hand, attains 81.7% and 79.6% F1-scores for these datasets. This clearly shows the improvement over the baseline and the benefit of the optimization approach. As *N* is increased both approaches’ performance deteriorates as expected, but the gap between the two approaches stays the same. For *N* = 50, the optimization-based approach achieves an F1-score of 70% and 67%, for OpenSNP and HapMap datasets, respectively. This is a typical beacon size considered in the literature (7; 8; 28; 16). A 70% F1-score represents a strong balance between precision and recall, especially in a realistic, challenging and imbalanced task as this. This serves as a meaningful improvement over the baseline performance and shows the feasibility of such a beacon reconstruction attack. Note that the correlation matrix *C* is estimated from the OpenSNP dataset and the results obtained on the HapMap dataset with this correlation matrix show that even if the source of the correlations is not from the same population and is obtained from a rather noisy community-driven public dataset, the optimization-based reconstruction method can achieve high performance. On the other hand, performance decrease when *N* doubles from 50 to 100 is only 4% and the gap between the optimization-based approach and the baseline is preserved. This indicates that the method can scale well while the problem size is doubled. For the same experiment with varying SNP set sizes can be found in the Supplementary Figures 1 and 2, for OpenSNP and HapMap datasets, respectively. We observe the same trend in both figures.

**Fig. 1.**
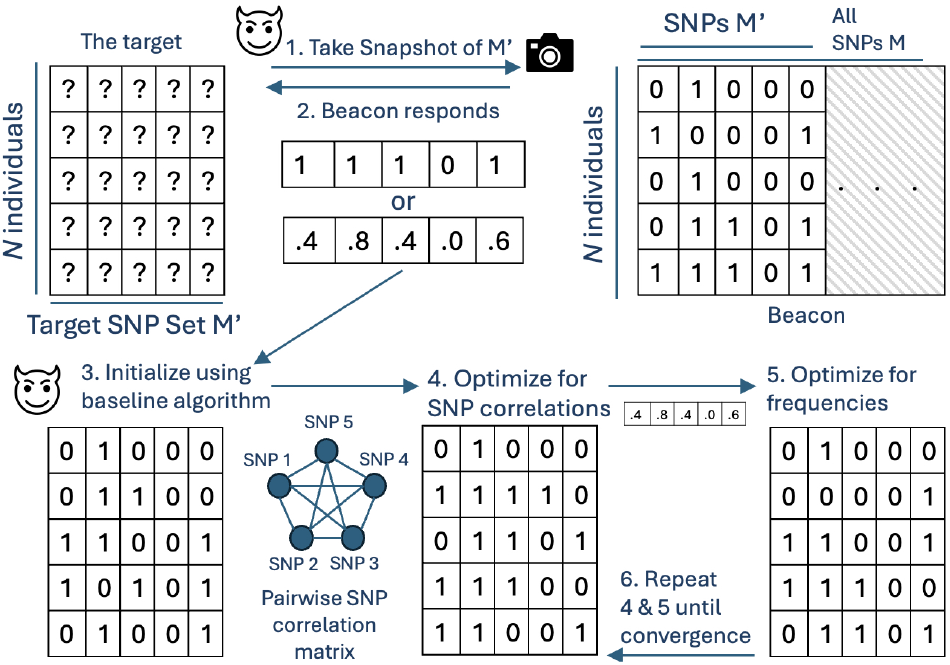
The system model of our approach. The attacker finds out *N* and decides on the subset of SNPs *M*^′^ to reconstruct. 1. The attacker takes the snapshot of the beacon for *M*^′^. 2. Beacon can respond with allele frequencies or yes/no for each SNP. In the latter, the attacker estimates allee frequencies from population. 3. The attacker initializes the matrix using baseline algorithm, and in 4., updates the assignments such that the SNP correlations are preserved. 5. The attacker optimizes for the original allele frequencies obtained. 6. Steps 4 and 5 are alternated until convergence.

**Fig. 2.**
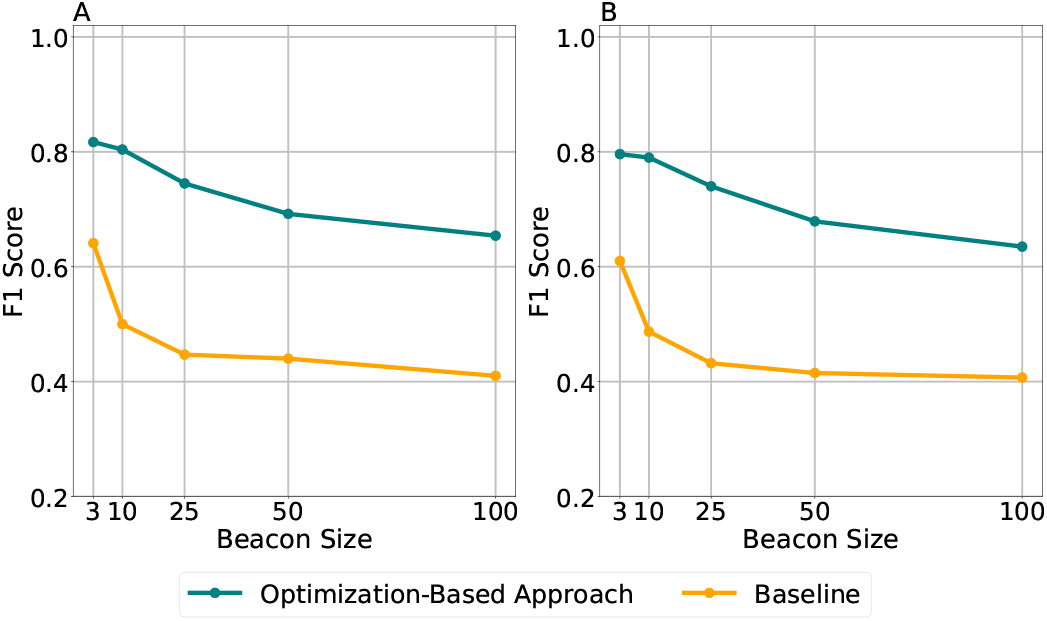
F1-Score comparison for |*M*^′^| = 1000 for varying beacon size (*N*). Plot A represents the reconstruction of the OpenSNP-based beacons, Plot B represents the reconstructions of the HapMap-based beacons.

#### Effect of Number of SNPs Targeted

Here, we evaluate the effect of varying the number of SNPs considered (|*M* ^′^|). First, the results in Supplementary Figure 1 and 2 demonstrate that the optimization-based approach performs substantially better than the baseline method for all |*M* ^′^| values, for both datasets. For example, in panel B of Supplementary Figure 1 where |*M* ^′^| = 50, the optimization-based method achieves an F1-score of 94% for 3 individuals in the OpenSNP dataset. In comparison, the baseline approach achieves 71% which assumes independence between SNPs. The performance gap arises due to the baseline’s tendency to make false assignments due to its inability to account for correlations between SNPs causing incorrect predictions of alleles. As |*M* ^′^| increases, both approaches’ performances go down, which is expected due to the increasing search space and complexity of the problem. However, we observe that with even 2000 SNPs considered and 50 individuals in the beacon (Supplementary Figure 1E), the performance of the optimization-based approach is 67%, which is a strong F1-score considering the size of the problem. The F1-score gap between predicting the assignment of 1000 SNPs (Figure 2) and 2000 SNPs (Supplementary Figure 1E) for 50 individuals is just 2.2%. This indicates that the model scales well even if the number of SNPs doubles. We observe a very similar pattern in the results obtained on the HapMap dataset as shown in Supplementary Figure 2.

#### Effect of already reidentified individuals

Here, we investigate the effect of the attacker having access to the genomes of a percentage of the beacon participants through data leaks and reidentification attacks as discussed in Section 3.4.3. We randomly select *p* = 20% and 40% of the individuals in both data sets and assume that their genomes are leaked to the attacker who has reidentified them as participants in the beacon dataset. The attacker fixes these individuals in **B**^′^ and reconstructs the rest (1 *− p*) of the individuals for varying numbers of SNPs |*M* ^′^| and beacon sizes *N* using the optimization-based method. To observe the effect of individuals already reidentified in reconstruction performance, we reconstruct the remaining (1 *− p*) of the individuals in the beacon using the same method, but without the assumption of the attacker having access to the genomes of any person in the beacon and compare the results. This is indicated as “Without known *p*” in Figure 3.

**Fig. 3.**
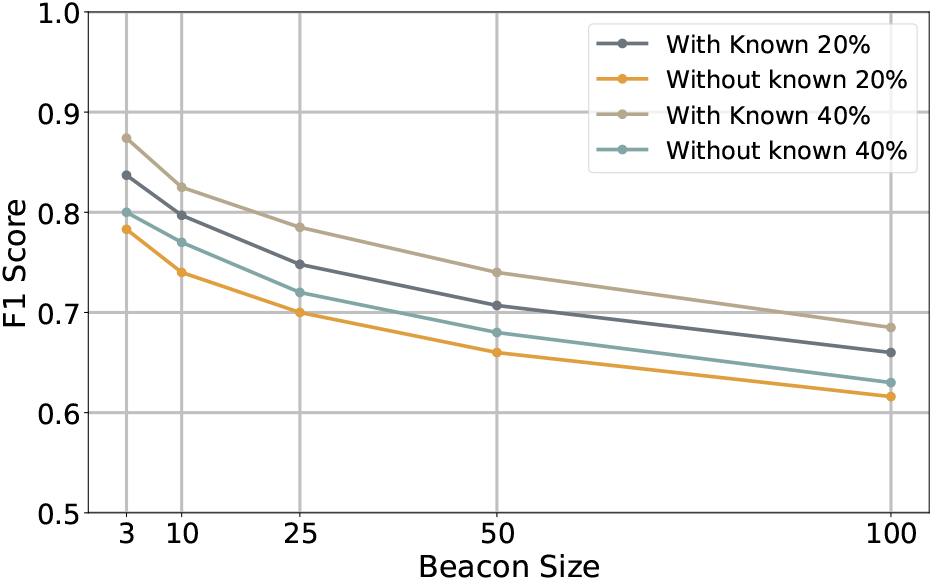
F1-Score comparison for |*M*^′^| = 1000 in the HapMap dataset for varying beacon size (*N*). The results compare the performance of the attacker to reconstruct same number of genomes with access to genomes of *p* = 20% (“With Known 20%”) and *p* = 40% (“With Known 40%”) of participants, and without this information (“Without known 20%” and “Without known 40%”).

We observe that having access to a portion of the beacon leads to higher performance for both *p* = 20% and 40% compared to reconstructing the same number of individuals but without using the information of known 20% and 40%. The improvement in F1-score is on average is 5.59%. The larger *p* leads to better reconstruction performance as the search space is smaller for the optimizer. The F1-score gap between *p* = 20% and 40% is on average is 3.2%. For a beacon with 50 individuals, if the attacker has access to only 10 genomes, the reconstruction achieves an F1-score of 70.7% which makes the attack even more stronger, further underscoring the importance of the threat. For the same experiment with varying SNP set sizes can be found in the Supplementary Figure 3 for the HapMap dataset where we observe the same trend.

#### Effect of Beacons Reporting Allele Frequency

All results reported above, assume that the beacon responds to queries with AFs which is implemented in various beacons such as THL Biobank Beacon (32) and Progenetix Beacon+ (33). However, beacons can also respond by the presence or absence of the alleles instead of the frequency (i.e., binary). Clearly, this returns less information compared to AFs and makes it more difficult for the attacker to reconstruct the dataset. In this case, we assume that the attacker estimates the AFs in the beacon dataset by using background AFs observed in the population or in other public datasets and investigate the performance of the attack in this setting.

We use 64 left-out CEU HapMap samples to estimate the frequencies that would have been reported by the beacon interfacing the HapMap dataset (with varying *N* and |*M* ^′^|) instead of a “Yes/No” answer. That is, the attacker simulates the beacon and the underlying dataset using the left-out samples and uses the frequencies in the simulated dataset if they receive a “Yes” response from the beacon. If they receive a “No” response then the attacker sets the frequency for that SNP as zero. To consider the case where the attacker does not have access to the population AFs or the public AFs are not representing the beacon dataset well. For instance, the genetic background in the dataset can be mixed. To simulate this, we estimate the AFs using the Mexican population in the HapMap dataset as well.

The results are shown in the Figure 4. We observe that in the HapMap beacon with *N* = 50 and |*M* ^′^| = 1000, the optimization-based algorithm achieves an F1-score of 53.4%.

**Fig. 4.**
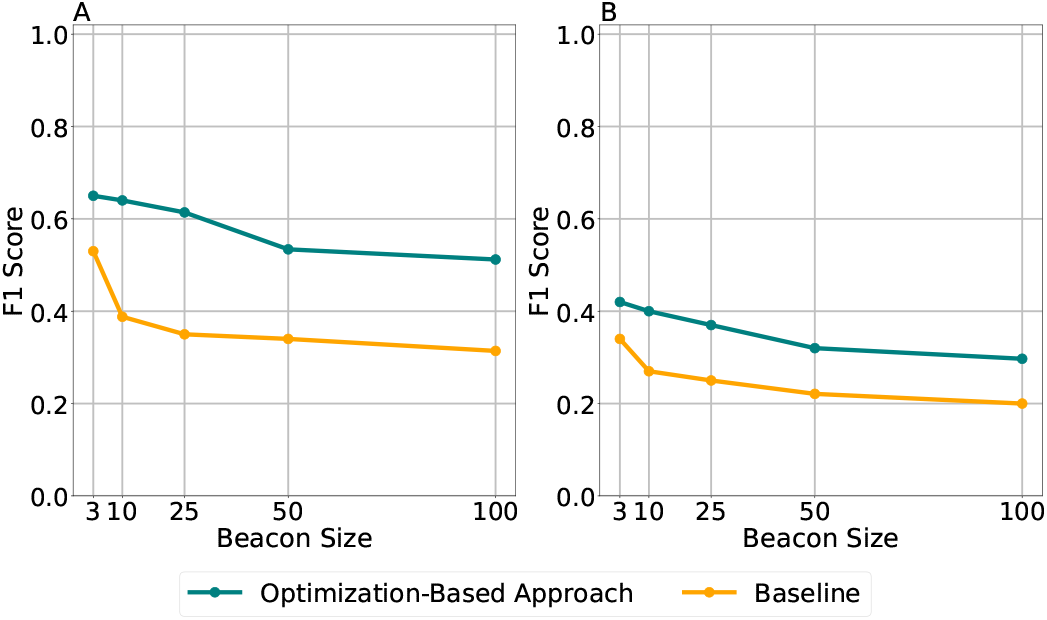
F1-Score comparison for |*M*^′^| = 1000 when frequency is unknown. Plot A represents reconstruction from 65 left-out samples of HapMap dataset, Plot B represents reconstruction using Mexican samples.

This is a decrease of 14% in the F1-score obtained when the beacon reports actual allele frequencies in the dataset.

Clearly exact AFs reported by the beacon result in more precise reconstruction and the algorithm loses power. However, the baseline algorithm performs much worse (37%). This indicates the model still remains effective in less ideal scenarios such as using background frequencies from the same population. For the same experiment with varying SNP set sizes, can be found in the Supplementary Figure 4 for the HapMap dataset.

With MAFs in the Mexican population, the attacker obtains an F1 score of 32%. For the same experiment with varying SNP set sizes, can be found in the Supplementary Figure 5 for the HapMap dataset. The results show that the use of AFs in the Mexican population substantially decreases performance. Note that this represents an extreme case where we estimate the frequencies using incorrect background information. However, these results show that beacons with genetically uniform content leak more information and are more prone to reconstruction attacks.

#### Time Performance

Supplementary Figure 6 shows the time requirement to reconstruct the beacons with varying *N* and |*M* ^′^|. The optimization-based algorithm scales linearly with *N* and |*M* ^′^|. With *N* = 100 and |*M* ^′^| = 2000, the attack takes around 10 hours which was the most time-consuming case in our analyses.

The attack is performed offline and the results show that it can scale for large numbers of variants and beacon sizes.

## Discussion and Conclusion

In this work, we introduced the first-of-its-kind, beacon reconstruction attack which can reconstruct large numbers of variants in all beacon participants’ genomes with high performance. We tested the limits of the approach under various varying factors such as beacon size, number of target variants, percentage of already identified individuals in the beacon and assumptions on summary statistics. Our results show that the attack is feasible. Required time scales linearly with increasing number of participants and targeted variants. The results decline again linearly with increasing number of participants but with a shallow slope. We have tested the attack in a challenging setting where the SNP correlations are estimated from the OpenSNP dataset which does not clearly reflect the actual correlations in the CEU population. Yet, our attack was effective.

The beacon protocol allows AFs to be reported (5) and many beacons implement this option. We clearly show that if beacons report allele frequencies, the beacon reconstruction is more effective compared to simple “Yes/No” responses. The attacker can estimate the frequencies using background AFs and the attack is still feasible but is less powerful. Therefore, suggest administrators implementing the beacon protocol to avoid AF reporting option for safety. In this case, we also show that the attack is again more effective if the dataset contains individuals from a single population. Thus, larger number of individuals with diverse genetic backgrounds can help mitigate such beacon reconstruction attacks.

Our suggested approach relies on pairwise correlations among variants. Yet, other approaches suggest using higher-order correlations can provide more information and can be used to reconstruct other parts of the genomes (9; 22). While we relied on a randomly selected subset of variants to test the limits of the model, a more carefully selected subset of variants can be targeted using our approach to form a baseline. This can be followed by an independent second stage where the missing parts of the genomes of the victims can be further reconstructed using higher-order correlations as done in (22). This means the beacon reconstruction attack which focus on pairwise correlations can be paired with a second attack. This follow-up attack can further reconstruct the remaining missing parts using higher-order correlations independently from the beacon summary statistics to uncover more variants. We will consider this in the future work.

One can also consider selecting a relatively more correlated subset of variants. We have investigated this issue and reconstructed subsets of SNPs with low, moderate and high average correlations. Yet, the performance of the reconstruction was not substantially different indicating the method is robust against this effect as long as it can rely on confidently estimated correlation matrix. In our experiments, we used a correlation matrix estimated from the OpenSNP dataset to reconstruct the a beacon simulated using individuals from the CEU population. More confident estimation of the correlation matrix can yield stronger attacks, indicating that there is still room for improvement.

## Acknowledgements

We thank Göktuğ Gürbüztürk, Efe Erkan, Deniz Aydemir, İrem Aydın, Kerem Ayöz, and Erman Ayday for their contributions to this study.

## Footnotes

1 https://ftp.ncbi.nlm.nih.gov/hapmap/genotypes/2010-08/_phaseII+III/forward/

