## Supplementary Material for "Beacon Reconstruction Attack: Reconstruction of genomes in genomic data-sharing beacons using summary statistics"

### 1 Supplementary Figures

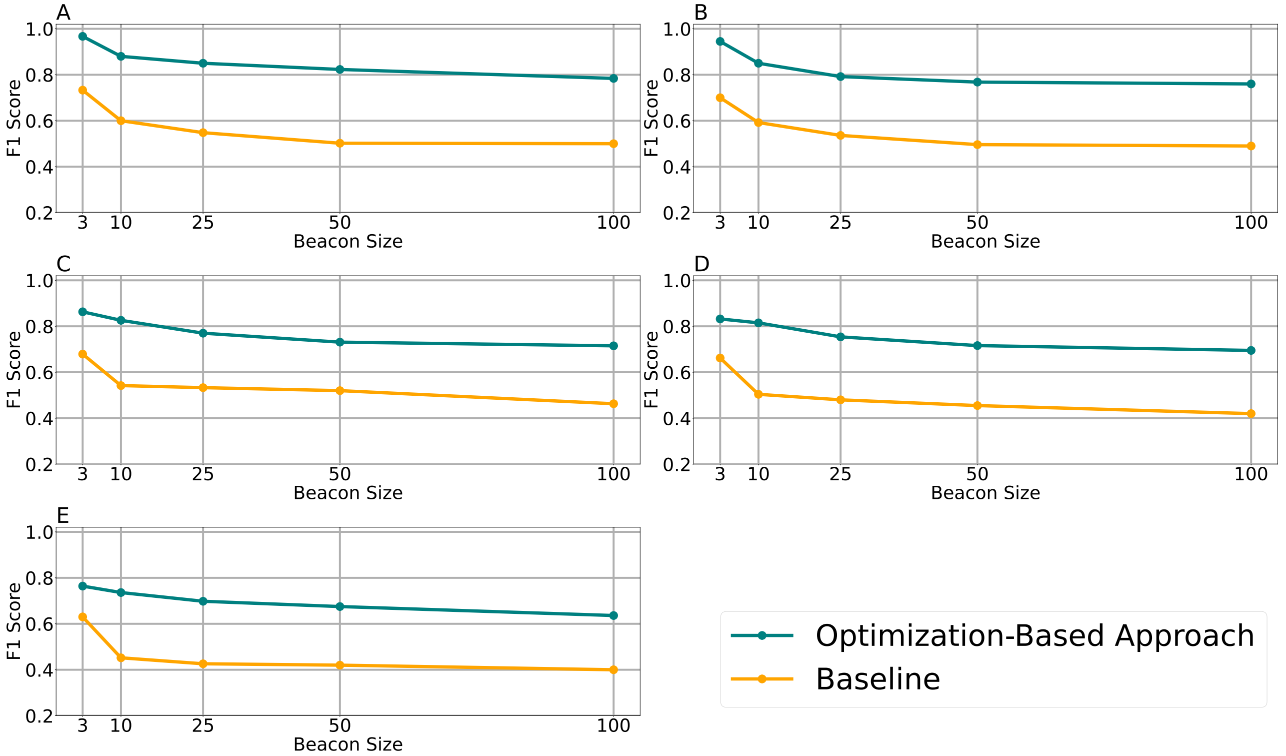


Supplementary Figure 1: F1-Score comparison across different ($|M'|$ = 30, 50, 100, 500, 2000) and varying numbers of individuals in the OpenSNP dataset. Plots A–E correspond to these $|M'|$, respectively.

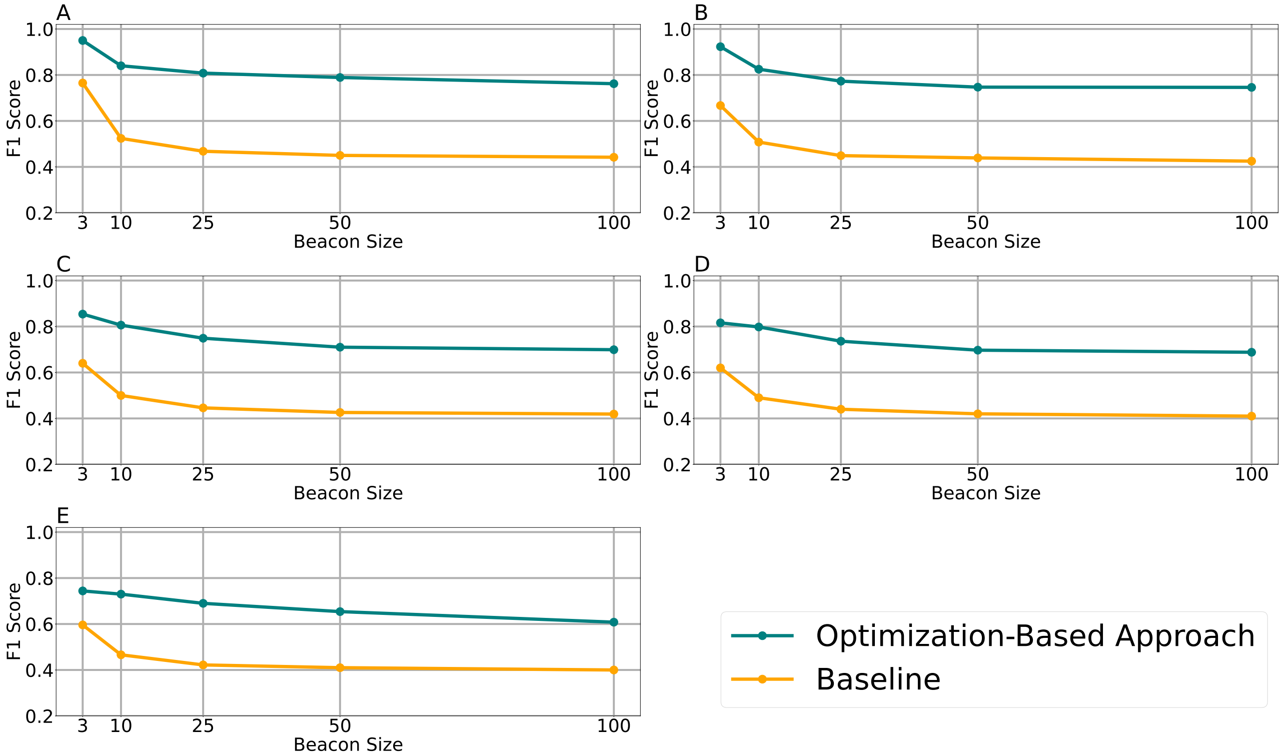


Supplementary Figure 2: F1-Score comparison across different ($|M'|$ = 30, 50, 100, 500, 2000) and varying numbers of individuals in the HapMap dataset. Plots A–E correspond to these $|M'|$, respectively.

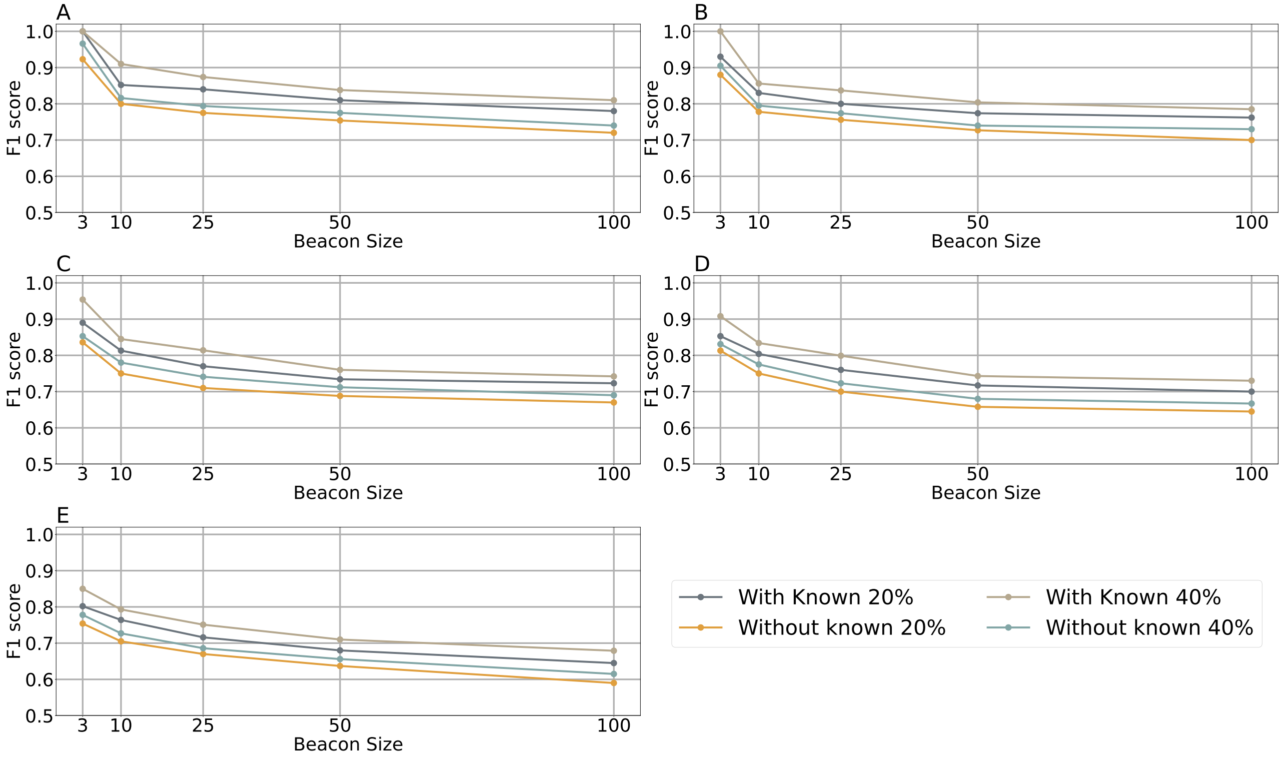


Supplementary Figure 3: F1-Score comparison across different ($|M'|$ = 30, 50, 100, 500, 2000) and varying numbers of individuals in the HapMap dataset. Plots A–E correspond to these $|M'|$, respectively. The results compare the performance of the attacker with access to genomes of $p$ = 20% ("With Known 20%") and $p$ = 40% ("With Known 40%") of participants, and without this information ("Without known 20%" and "Without known 40%").

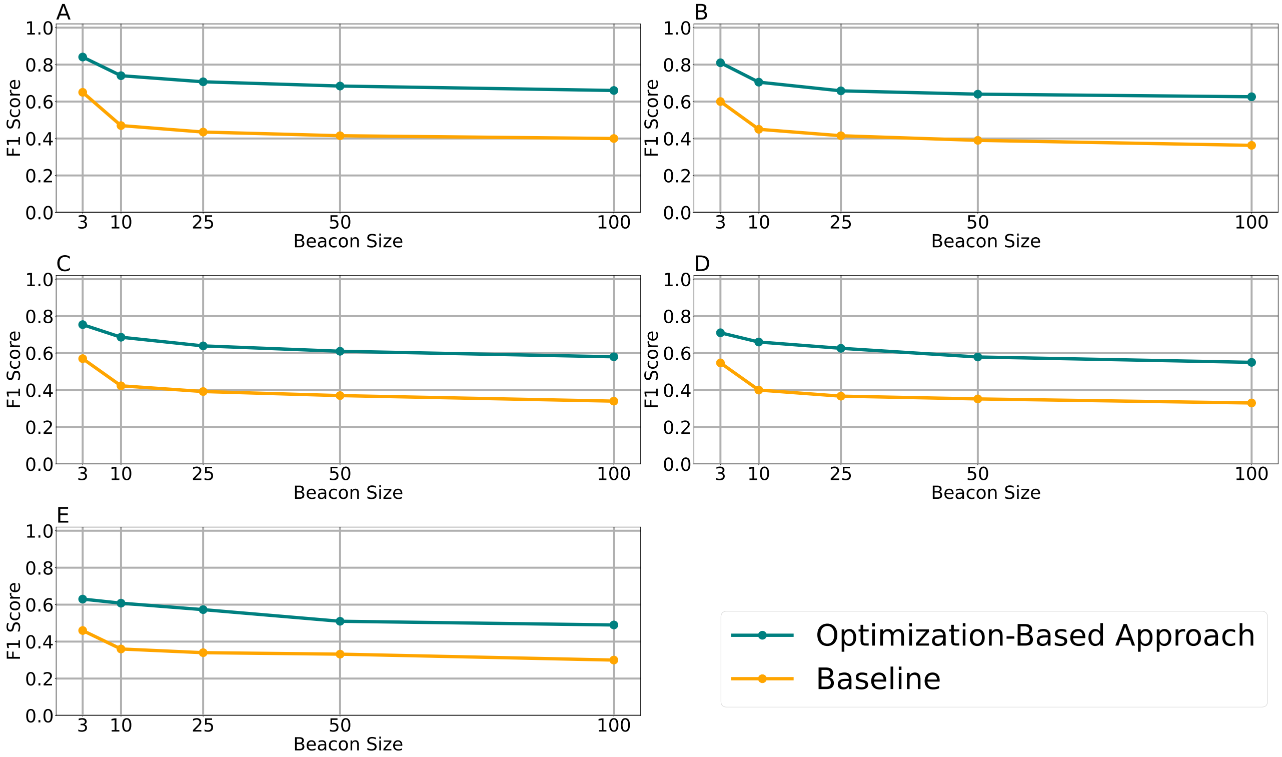


Supplementary Figure 4: F1-Score comparison using MAFs of 64 left-out individuals out of 164 of HapMap dataset. The comparison is across different ($|M'| = 30, 50, 100, 500, 2000$) and varying numbers of individuals. Plots A–E correspond to these $|M'|$, respectively.

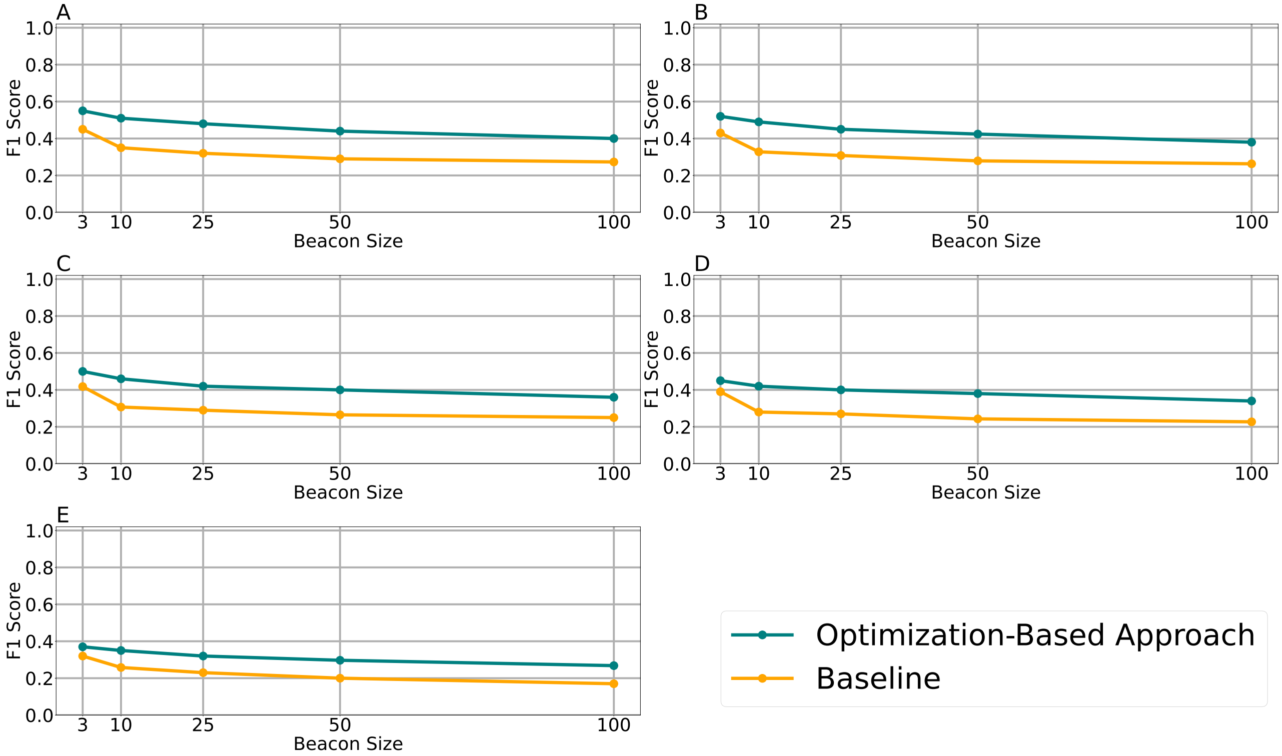


Supplementary Figure 5: F1-Score comparison for the reconstruction of the HapMap dataset using Mexican population MAFs, evaluated across varying numbers of individuals and ($|M'|$ = 30, 50, 100, 500, 2000). Plots A–E correspond to these ($|M'|$, respectively.

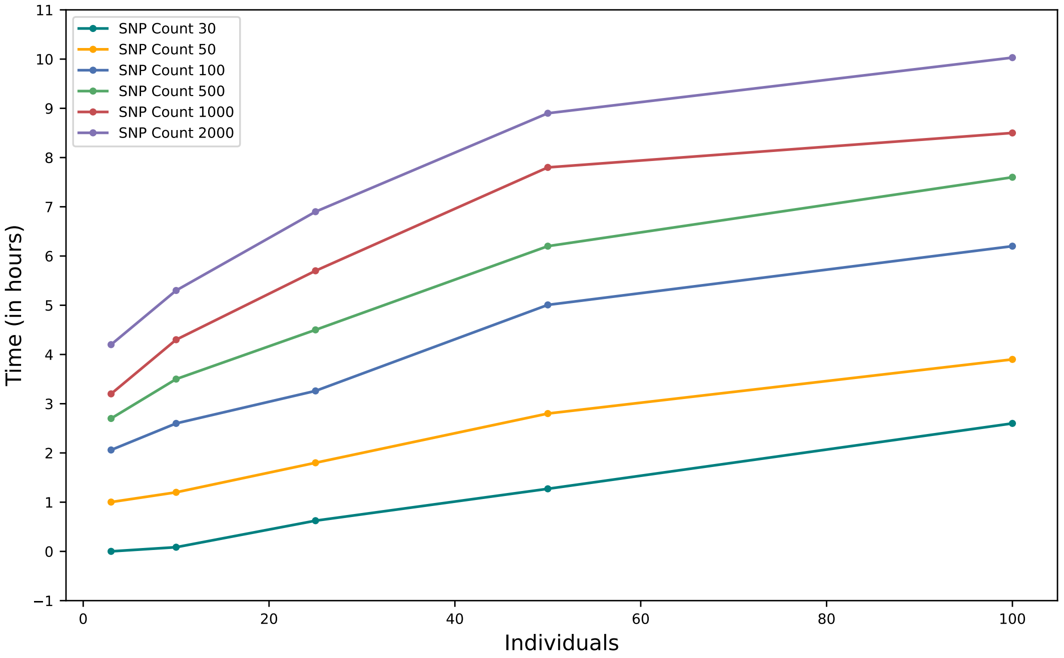


Supplementary Figure 6: Time Analysis Graph across ($|M'|$ = 30, 50, 100, 500, 2000) and varying numbers of individuals
